# Mutational scans reveal differential evolvability of *Drosophila* promoters and enhancers

**DOI:** 10.1101/2022.10.17.512533

**Authors:** Xueying C. Li, Timothy Fuqua, Maria Elize van Breugel, Justin Crocker

## Abstract

Rapid enhancer and slow promoter evolution have been demonstrated through comparative genomics. However, it is not clear how this information is encoded genetically and if this can be used to place evolution in a predictive context. Part of the challenge is that our understanding of the potential for regulatory evolution is biased primarily toward natural variation or limited experimental perturbations. Here, to explore the evolutionary capacity of promoter variation, we surveyed an unbiased mutation library for three promoters in *Drosophila melanogaster*. We found that mutations in promoters had limited to no effect on spatial patterns of gene expression. Compared to developmental enhancers, promoters are more robust to mutations and have more access to mutations that can increase gene expression, suggesting that their low activity might be a result of selection. Consistent with these observations, increasing the promoter activity at the endogenous locus of *shavenbaby* led to increased transcription yet limited phenotypic changes. Taken together, developmental promoters may encode robust transcriptional outputs allowing evolvability through the integration of diverse developmental enhancers.

**Quote:** “Regulators, mount up [at transcriptional promoters].” - Warren G & Nate Dogg, 1994

## Introduction

Mutations may be largely random, but the loci of evolution are not. Through analyzing causal variants underlying natural variation, previous studies have found that specific genes or nucleotide substitutions are more often used in evolution than others (Stern and Orgogozo 2008; Chan et al. 2010; Martin and Orgogozo 2013). For example, *cis*-regulatory changes are shown to be favored by long-term evolution and morphological traits (Stern and Orgogozo 2008), although their relative contribution to evolution compared to coding changes is still under debate (Hoekstra and Coyne 2007; Stern and Orgogozo 2009). Therefore, evolution may be predictable if we gain a more comprehensive understanding of the roles of different kinds of molecular changes and their possible contributions to evolution. However, the ability to predict evolution requires a full construction of the genotype-to-fitness map, which is difficult to achieve by analyzing naturally occurring variations that are limited in number and shaped by selection (Perkins et al. 2022).

Mutational scans of regulatory and coding sequences in microorganisms and cell lines have begun to map genotype-to-phenotype relationships in broad sequence spaces (Patwardhan et al. 2012; Li et al. 2016; Venkataram et al. 2016; Kinsler et al. 2020) and to reveal principles of regulatory grammar (Sharon et al. 2012; Kwasnieski et al. 2012; Qi et al. 2022) and adaptation (Metzger et al. 2015; Park et al. 2022). However, such data have been largely lacking for developmental systems, where the additional challenge is to understand how mutations impact the spatial and temporal pattern of gene expression across development and populations. Recently, mutational scans have been applied to developmental enhancers in fruit flies, where it was found that almost all mutations altered gene expression (Fuqua et al. 2020). This study was further extended to additional elements (Galupa et al. 2023), suggesting that developmental enhancers are often sensitive to perturbation, and may be highly constrained.

Metazoan promoters are traditionally thought to be functionally separated from enhancers, with the former primarily interacting with the transcription machinery (e.g. Pol II) and the latter interacting with transcription factors carrying spatial and temporal information. However, recent studies suggest that the boundary between promoters and enhancers can be blurry: enhancers can initiate certain levels of transcription, just like promoters, and many known promoters can influence transcription initiation of other genes, which is the classical definition of enhancers (Haberle and Stark 2018; Andersson and Sandelin 2020; Ramalingam et al. 2022). From an evolutionary standpoint, it has been found that rapid evolution of enhancers is a general feature of mammalian genomes (Villar et al. 2015). In contrast, the genomic enrichment of key histone marks H3K27 acetylation and H3K4 trimethylation associated with promoters was partially or fully conserved across these species, suggesting that there is slow evolution of promoters in animal genomes. However, it is not known how this information is encoded genetically at developmental promoters, which have been distinguished from “housekeeping” promoters by their distinct properties in the epigenetic and sequence signatures (Lenhard et al. 2012), the level of PolII stalling (Zeitlinger et al. 2007) and enhancer preferences (Zabidi et al. 2015).

Here, to understand if mutations in developmental promoters have a different distribution of effects on gene expression from those in enhancers, we examined random mutation libraries of three *Drosophila* promoters and compared them to a previously surveyed *Drosophila E3N* enhancer. In contrast with the previous findings that enhancers may be highly sensitive to mutations (Fuqua et al. 2020), we found that mutations in these promoters sometimes change the level of gene expression, but never the spatial pattern of expression. Together, these findings suggest that developmental promoters may encode robust transcriptional outputs allowing evolvability through the integration of diverse developmental enhancers.

## Results

We focused our analyses on the regulatory sequences of *shavenbaby (svb)*, a gene that encodes an essential regulator of trichome development in *Drosophila*. The evolution of the *svb* regulatory regions has been extensively studied due to contributions to phenotyic evolution across many *Drosophila* species (Sucena and Stern 2000; Frankel et al. 2011; Crocker et al. 2015; Preger-Ben Noon et al. 2016). Through these works, seven transcriptional enhancers have been characterized for *svb*; each integrates information from multiple patterning networks giving rise to the overall expression of *svb* across the embryo (**Fig. 1A**).

**Fig. 1.**
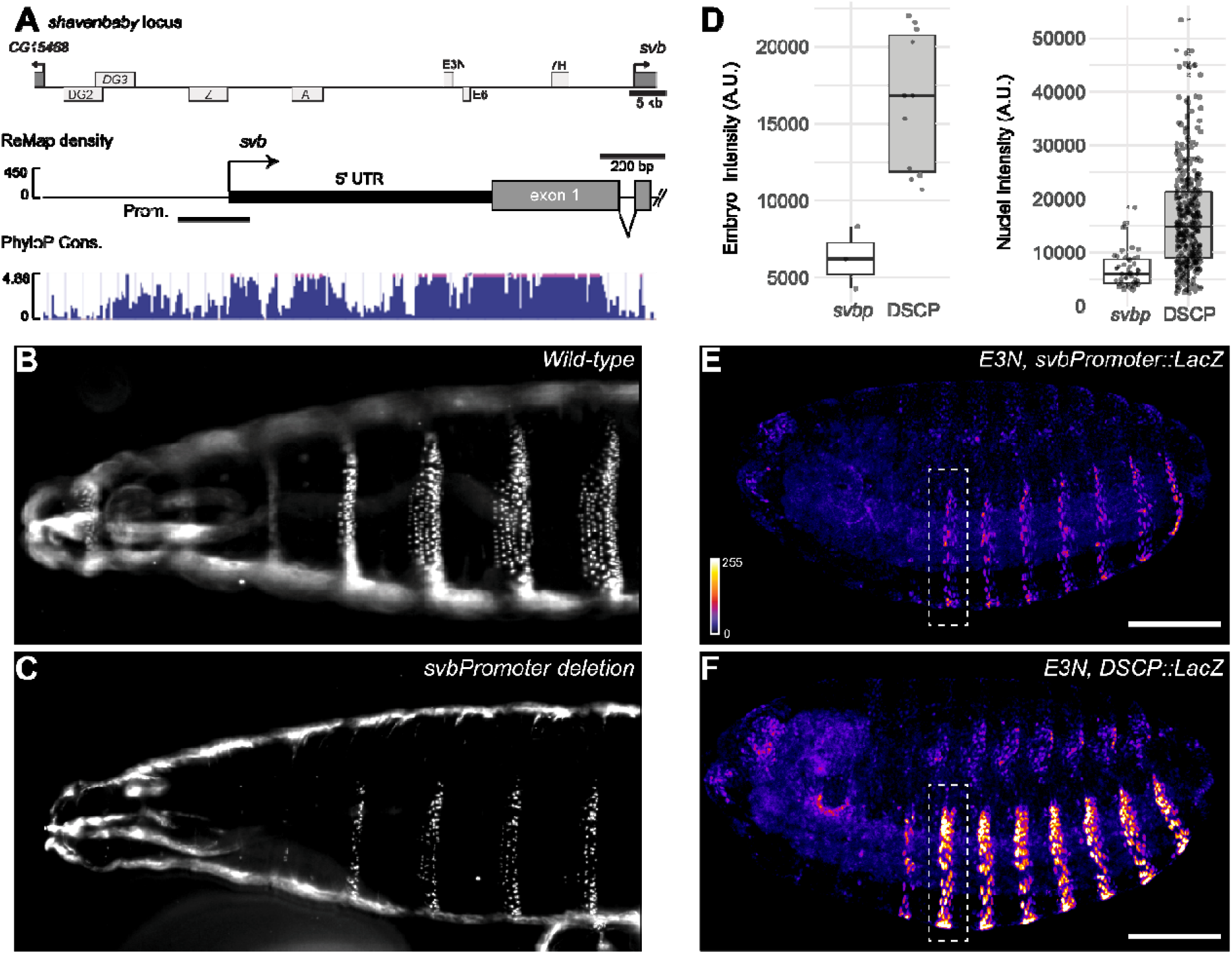
*LacZ* expression driven by *svb* promoter and *DSCP*. (**A**) *shavenbaby* locus. The regulatory region of *svb* spans ∼87kb, consisting of seven enhancer regions (top) (Stern and Frankel 2013). The promoter of *svb* is 226bp-long, with high regulatory activity (ReMap score, middle) and conservation level (PhyloP score of 124 insects, bottom). (**B, C**) Cuticles of wild-type and *svbp* deletion flies, respectively. (**D**) Levels of *lacZ* expression driven by *svb* promoter (*svbp*) and DSCP under the control of *E3N*. Nuclei intensity was quantified by extracting average intensity around local maxima of A2 stripe (shown by white boxes in **E** and **F**). Left, mean intensity per embryo, p < 0.01. Right, nuclei intensity across embryos, p < 0.001. A. U., arbitrary unit. P values were from Wilcoxon tests. (**E, F**) Representative images of stage15 embryos, showing the pattern of *LacZ* expression in abdominal stripes driven by *E3N*-*svbp* and *E3N*-DSCP respectively, detected by anti-beta-Gal staining. Scale bar = 100 um.

To explore how the native *svb* promoter integrates these diverse activities, we tested the activity of the *svb* promoter (*svbp*) using integrated reporter gene assays. The *svbp* shows high regulatory activity based on ReMap density (Hammal et al. 2022) and is conserved among *Drosophila* species (Siepel et al. 2005) (**Fig. 1A**). It does not contain TATA-box or other strong transcription motifs, consistent with signatures of developmentally regulated promoters in *Drosophila* (Lenhard et al. 2012). Deletion of *svbp* resulted in severe depletion of ventral trichomes (**Fig. 1B-C**), recapitulating *svb* mutant phenotypes (Payre et al. 1999), although these lines were homozygous viable. In order to understand how different promoters control levels and patterns of transcription activities driven by developmental enhancers, we generated reporter constructs of *svb* promoter and *Drosophila* synthetic core promoter (DSCP), an artificially engineered promoter known to drive high levels of expression (Pfeiffer et al. 2008). Both promoters were placed downstream of the *E3N* enhancer of *svb*, which drives expression in a pattern of eight stripes in the abdominal region (A1 to A8) in stage 15 embryos (Fuqua et al. 2020). The design with the *E3N* enhancer was necessary because, in the absence of an enhancer, the construct (with *hsp70* promoter) only drove a low level of background expression in stage 15 embryos (Galupa et al. 2023). We found that the two promoters drove different levels of reporter gene expression in the stripes, using the second abdominal stripe (A2) as a focal region for quantification (**Fig. 1D, Fig. S1**). The nuclei intensity from *DSCP* was on average 2.4-fold higher than that of *svbp*. However, we found no differences in the overall gene expression patterns in the stripes (**Fig. 1E-F**). Additionally, we tested constructs without a promoter and with a random sequence as the promoter. We found that the reporter expression was below the level of detection in both lines (**Fig. S2**), ruling out any promoter activity contributed by *E3N* in the constructs.

In order to understand the evolutionary potential of promoters in regulating the level and pattern of gene expression in a developmental context, we generated random mutation libraries of *svbp* and *DSCP* at a mutation rate of 1-2%, in a similar manner to our previous mutational scan on the *E3N* enhancer (Fuqua et al. 2020). In the previous study, which examined the mutational profile of *E3N* in combination with the *hsp70* promoter, single point mutations in *E3N* almost always decreased the gene expression level (18/18 variants). They often changed the state or locations of gene expression (11/18, 61%). In the follow-up studies which examined *E3N* variants with 1-10 mutations, it was found that the majority of variants caused decreases in the level of gene expression (83/91 based on median intensity, 91%) (Galupa et al. 2023) and reduction in the number of nuclei in A1 to A8 stripes (81/91, 89%) (Fuqua 2021). We independently recapitulated these results by analyzing ten randomly selected lines with 2-3 mutations from the *E3N* library. We found that all ten lines had a reduced number of nuclei expressing *lacZ* (**Fig. 2A-C, Fig. S3**) and that seven showed reduced levels of expression (FDR-adjusted p < 0.05, Wilcoxon test, **Fig. 2A**). Together, these results are consistent with the previous finding that enhancers encode dense spatial information (Fuqua et al. 2020; Le Poul et al. 2020; Galupa et al. 2023).

**Fig. 2.**
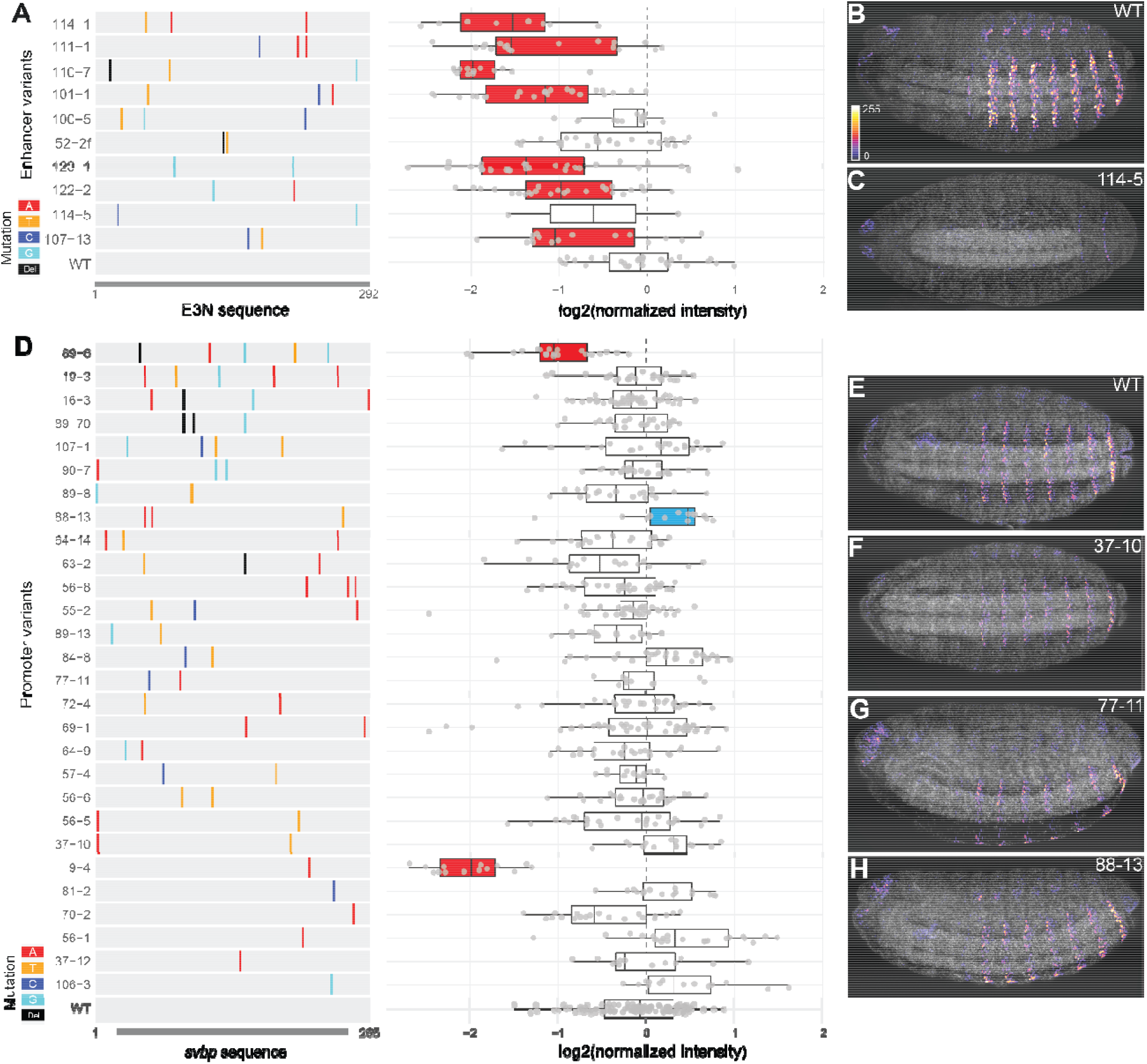
Different mutational profiles of *svb* enhancer and promoter. (**A**) Mutations and expression level of *E3N* variants, with representative images in (**B**-**C**). (**D**) Mutations and expression level of *svb* promoter variants, with representative images in (**E**-**H**). The mutant lines were ordered by the number of mutations, from low (bottom) to high (top). Colored lines show the position and identity of mutations. Del, deletion. The level of expression was represented by mean intensity of nuclei in the A2 stripe in each embryo, normalized to internal wild-type controls within each batch. The color of boxplots indicates significant difference from wild-type, tested by Wilcoxon test within batches (FDR-adjusted p < 0.05): red, reduced expression. Blue, increased expression. The grey channel in (**B-C**) and (**E-H**) shows DAPI staining.

In contrast, we analyzed 28 *svb* promoter variants (**Fig. 2D-H**), together covering 58 base pairs, and did not find any variants changing the pattern of gene expression (representative images in **Fig. 2F-H**). Unlike the enhancer, only two variants showed significantly lower expression than the wild-type *svb* promoter (**Fig. 2D**). Furthermore, one line showed higher expression levels than the wild-type promoter. The three lines with a changed expression level (89-6, 88-13 and 9-4) together contained seven unique mutations, and three were at highly conserved positions at the proximal end of the promoter (**Fig. S4A**), suggesting potential functional relevance. However, we did not find a significant correlation between the presence of phenotypic effect and the level of conservation [phyloP score from 124 insects (Siepel et al. 2005)], possibly due to the small sample size (Wilcoxon test, p > 0.05). Interestingly, line 9-4 harvested a single T-to-A mutation at the first nucleotide position of the *svb* 5’ UTR, which showed the most severe reduction in the level of gene expression in the library (**Fig. 2D**), suggesting a potentially critical role of transcription start sites. Taken together, given that the promoter library had a comparable mutation rate to the *E3N* enhancer library (on average 1%), our results suggest that developmental promoters are more robust than enhancers when subjected to the same mutation load.

We next extended our analysis to the *Drosophila* synthetic core promoter (DSCP). The DSCP was created by adding initiator (Inr) motif, motif ten element (MTE) and downstream promoter element (DPE) to a TATA-containing promoter of the developmental gene *even skipped* (*eve*) — creating one of the strongest promoters available in fruit flies (Pfeiffer et al. 2008). We quantitatively analyzed 45 variants of DSCP, with the number of mutations ranging from 1-8 across the 255bp-long sequence and an average mutation rate of 1.5% (**Fig. 3**). There were 117 nucleotide positions mutated in total, and 9 of them fell in the four functional motifs mentioned above (shaded regions in the left panel of **Fig. 3A**). We found that mutations in DSCP changed the expression level of the reporter gene more often than those in *svbp*, with 13 mutant lines showing significant changes, suggesting that the endogenous *svb* promoter might be more robust than the synthetic promoter. Among the variants showing changes in expression, 7-2 had a mutation in TATA, 17-2 had a mutation in Inr, and lines 13-11 and 49-1 both had mutations in MTE. However, we did not find a statistically significant enrichment of mutations in these transcriptional motifs in the lines that showed changes in expression vs. ones that did not (Fisher’s exact test, p > 0.05). Furthermore, there was not a correlation between the presence of an effect on gene expression and the level of conservation (for regions from *eve* promoter, **Fig. S4B**, Wilcoxon test, p > 0.05).

**Fig. 3.**
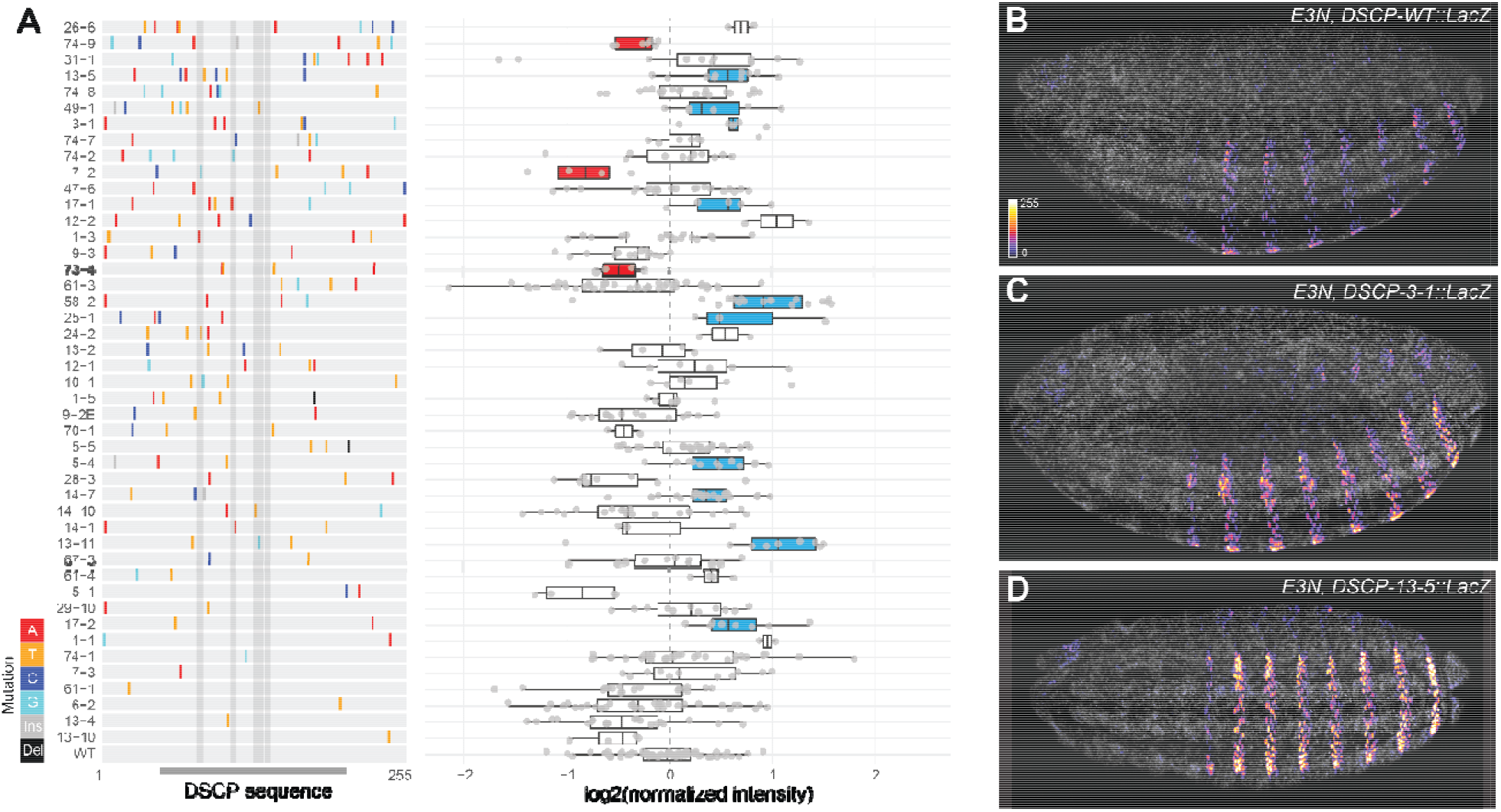
Mutational profiles of *Drosophila* synthetic core promoter. (**A**) Mutations and expression level of DSCP variants, with representative images in (**B**-**D**). The variants were ordered by the number of mutations, from low (bottom) to high (top). Grey shades show regions of TATA, Inr, and MTE-DPE motifs, respectively. Colored lines show the position and identity of mutations. Ins, insertion. Del, deletion. The level of expression was represented by mean intensity of nuclei in A2 stripes in each embryo, normalized to internal wild-type controls within each batch. The color of boxplots indicates significant difference from the wild-type promoter, tested by Wilcoxon test within batches on lines with a minimum sample size of three embryos (FDR-adjusted p < 0.05): red, reduced expression. Blue, increased expression. The grey channel in (**B-D**) shows DAPI staining. The images were background-subtracted and displayed in the same intensity range.

Interestingly, although DSCP drove a high level of expression, mutations in DSCP increased its activity even further in 10 mutant lines. It suggests that developmental promoters such as those of *svb* and *eve* might have the evolutionary potential to drive higher expression. However, it remains to be tested if strong transcriptional motifs such as those artificially engineered into DSCP are prerequisites of such evolvability. Due to the multiple mutational paths that led to higher levels of expression (either through transcriptional motifs or point mutations), developmental promoters might have been selected to maintain low transcriptional activity during evolution.

Consistent with our findings from the *svbp*, mutations in the DSCP did not change gene expression patterns, supported by the 45 lines quantified above (e.g. **Fig. 3B-D**) and 21 additional DSCP variants examined quantitatively (**Fig. S5A-C**). To further validate these results, we generated a mutation library of the *hsp70* promoter (*hsp70p*), a promoter commonly used to drive constitutive expression in *Drosophila* and used in the *E3N* library. Similarly, we did not find any variants causing a change in the expression pattern in the 31 variants examined (covering 74 out of 268 bp) (**Fig. S5D-F**). Together, these results are consistent with the traditional view of promoters encoding little spatial information (Serfling et al. 1985).

Although reporter constructs allowed us to examine the promoter variants in a controlled setting, it remains unknown whether the changes in transcription can lead to phenotypic outcomes at the endogenous locus, where complex promoter-enhancer interactions are involved. Therefore, we next tested if a change in the promoter activity at the endogenous locus could lead to phenotypic outcomes. We knocked out the *svb* promoter at its endogenous locus and replaced it with DSCP using CRISPR/Cas9. We found that the stronger DSCP promoter produced higher levels of transcription based on the local levels of nascent *svb* transcription compared to the endogenous promoter (**Fig. 4A-C**), consistent with the finding from reporter constructs. However, the changes in transcription levels did not directly translate into morphological changes, i.e., the pattern of ventral trichomes in larval cuticles (**Fig. 4D-E**): the DSCP knock-in rescued the knock-out phenotype (severe depletion of trichomes, **Fig. 1**) to the wild-type level, but there was no apparent differences in the trichome patterns from the wild-type.

**Fig. 4.**
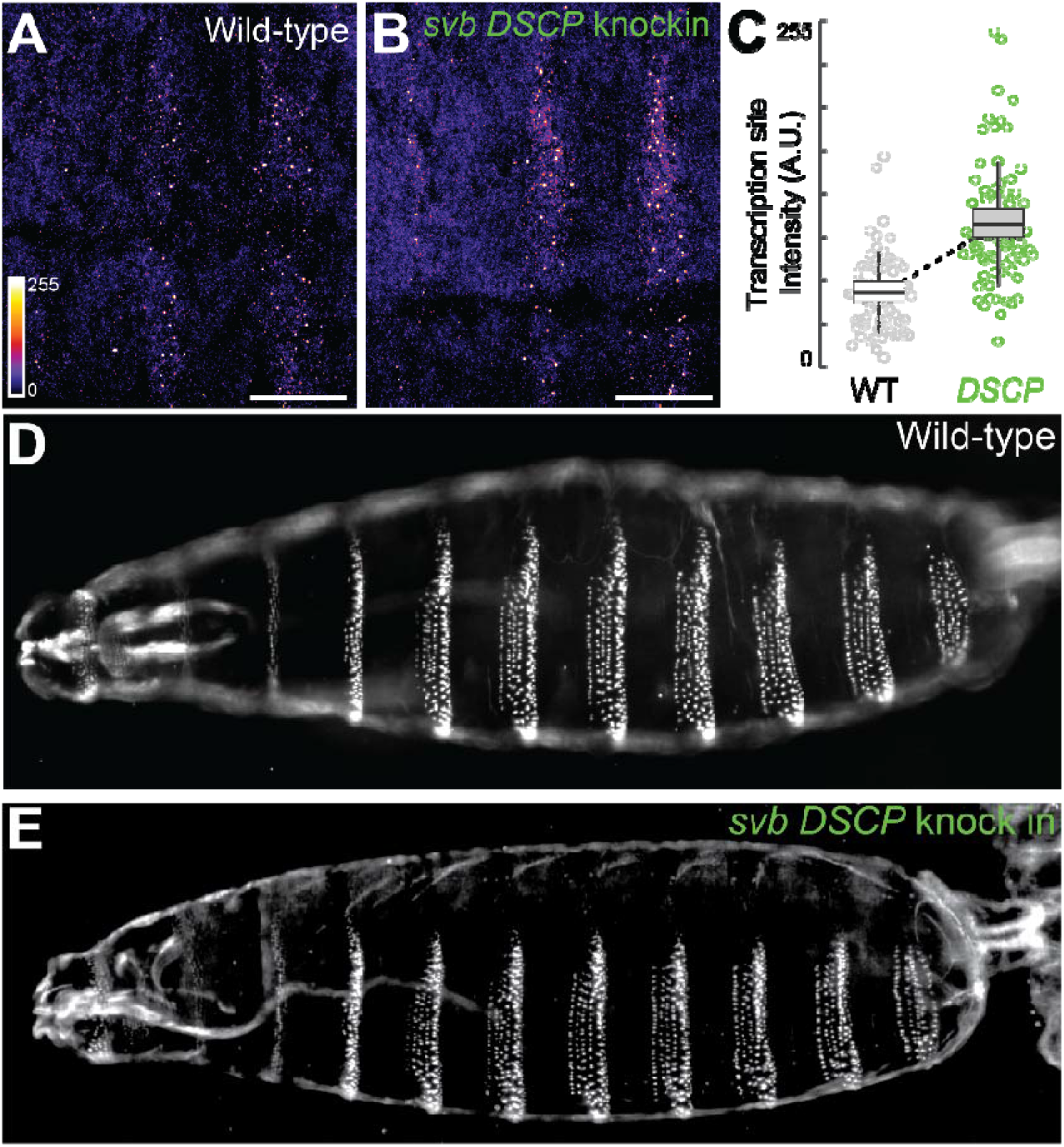
Replacing *svb* promoter with DSCP at its endogenous locus. (**A, B**) Transcription sites of *svb* in stage 15 embryos, detected by fluorescent in situ hybridization. Scale bar = 20 um. (**C**) Transcription site intensity, quantified across 3 embryos for each genotype. A. U., arbitrary unit. (**D-E**) Cuticles of wild-type and *svbp*Δ::*DSCP* larvae, showing ventral trichomes.

## Discussion

Although it remains debatable whether enhancers and promoters are functionally different elements from a transcription perspective (Haberle and Stark 2018; Andersson and Sandelin 2020), there is evidence that they are under different selective pressures, and possible evolutionary constraints. For example, comparative studies have shown that enhancer sequences undergo rapid sequence divergence while maintaining their regulatory functions via binding site turnover, consistent with stabilizing selection (Ludwig et al. 2000; Arnold et al. 2014). Gains and losses of enhancers were also found to be frequent in different lineages (Arnold et al. 2014; Villar et al. 2015). Promoters have been shown to exhibit higher levels of sequence divergence than surrounding regions in insects, possibly associated with an increased mutation rate (Main et al. 2013). Changes in promoters tend to be neutral (Hoffman and Birney 2010), consistent with our findings. They have also been shown to evolve slower than enhancers in mammals (Villar et al. 2015). Still, the level of constraint on promoters can differ among different types of promoters (Carninci et al. 2006), with highly constrained promoters associated with developmental functions (Lindblad-Toh et al. 2011).

Empirical characterization of the mutational space of enhancers and promoters in a developmental context was only made possible recently through mutational scans (Fuqua et al. 2020) and automation of embryo handling (Fuqua et al. 2021). Recent mutational scans have found that developmental enhancers encode dense regulatory information and are strongly constrained (Fuqua et al. 2020; Le Poul et al. 2020; Galupa et al. 2023). In this study, we found that *Drosophila* promoters have different mutational profiles from enhancers. At a comparable mutation rate to the previously published *E3N* enhancer library (Fuqua et al. 2020), variants in our promoter libraries did not show any changes in the pattern of gene expression (**Fig. 2-3, Fig. S5**). In contrast, almost all mutant lines of *E3N* changed the pattern (**Fig. 2, Fig. S3**). Mutations in promoters can change the level of expression in either direction (**Fig. 2-3**), whereas mutations in enhancers tended to reduce expression (**Fig. 2**) (Fuqua et al. 2020; Galupa et al. 2023).

Together, these findings suggest that *Drosophila* promoters might be more robust to mutations than enhancers. Interestingly, this difference seems to exist in yeast promoters, if one considers a yeast promoter to be a mixture of enhancer (binding transcription factors) and promoter (initiating transcription) sequences: in the study of *TDH3* promoter, mutations in transcription factor binding sites (“enhancer”) greatly reduced transcription whereas other mutations only fine-tuned the level of expression (Metzger et al. 2015). Additionally, our results indicated that promoters might have little potential to evolve new spatial patterns of expression, consistent with a previous finding that promoters were less likely to be repurposed as enhancers than the other way around in mammalian evolution (Carelli et al. 2018). However, this observation remains to be tested with more promoters and beyond the context of reporter constructs. Furthermore, the effects of promoter variants on the level of gene expression did not correlate with the number or the location (e.g. in TATA or other motifs) of mutations (**Fig 2-3**), suggesting that regulatory information might be randomly distributed in these promoters and a saturated mutational scan might be required to fully decode the regulatory potential of promoter sequences.

When comparing *svb* promoter and DSCP, it is clear that the endogenous *svb* promoter had low activity (**Fig. 1**), consistent with previous views (Haberle and Stark 2018). The fact that both *svbp* and DSCP had access to mutations that can increase the expression level (**Fig. 2-3**) suggested that the low activity of endogenous promoters might be a result of selection. Furthermore, the high-activity, artificially engineered promoter was more “evolvable” (or “breakable”) in the sense that many mutations led to changes in the level of gene expression, whereas the low-activity, endogenous *svb* promoter was relatively robust to mutations, suggesting that developmental promoters might have evolved to encode robust transcriptional outputs. This robustness may facilitate evolvability through the rapid integration of developmental enhancers that drive cell-type specific expression patterns – including novel or coopted elements (**Fig. 5**).

**Fig. 5.**
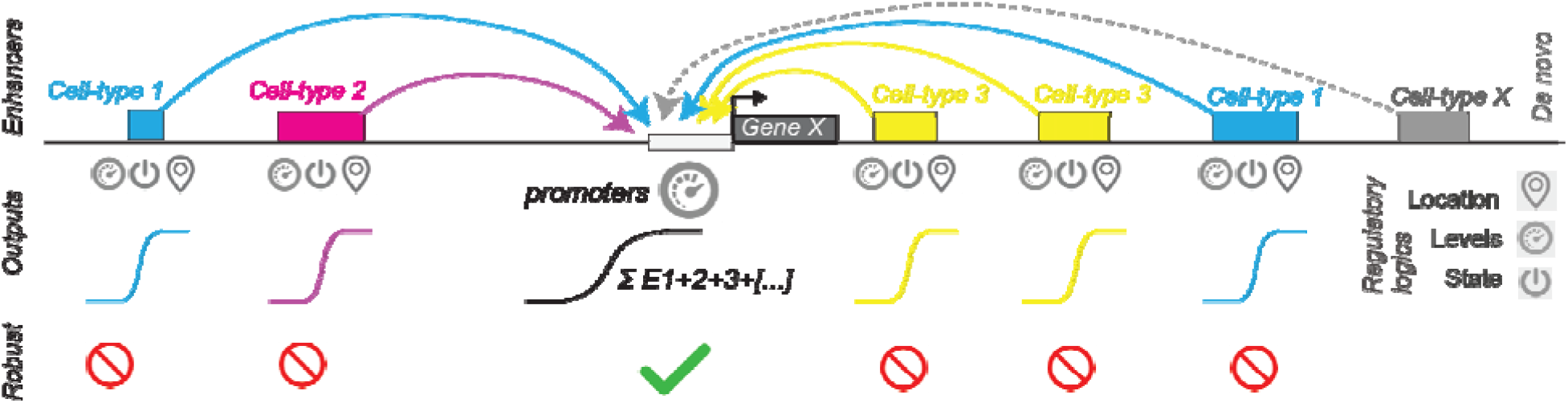
Model of enhancer and promoter evolution. Cell type-specific enhancers encode information for the location, levels and states of gene expression, whereas promoters encode information for the level of gene expression and integrate the transcriptional outputs from multiple enhancers. Promoters are relatively robust to mutations, allowing evolutionary changes through enhancers, including novel or coopted changes.

The relationship between the effects of *cis*-regulatory mutations on transcription and on fitness is often non-linear (Metzger et al. 2015; Bergen et al. 2016). In our study, we found that deletion of *svb* promoter led to a severe reduction of trichomes, similar to *svb* knockout phenotypes (Delon et al. 2003). However, changes in promoter activity at the endogenous locus of *svb* by knocking-in DSCP did not cause a change in larval trichome patterns (**Fig. 4**), suggesting that changes in transcription level could be buffered by the downstream network (Stern and Orgogozo 2008), consistent with developmental traits being highly robust systems (Siegal and Bergman 2002; Payne and Wagner 2015). Alternatively, the non-linear relationship could be explained by a threshold model, where the downstream patterning is elicited when *svb* transcription is above a certain threshold and a higher level of transcription does not change the patterning outcome (Delon et al. 2003).

Although promoters seem to be more robust to mutations than enhancers, the *svb* promoter shows a high level of sequence conservation, suggesting a certain degree of constraint. There could be a few explanations for this seemingly conflicted observation. First, promoters might be more “essential” to transcription than enhancers because transcription is usually closely associated with one promoter but possibly multiple enhancers with redundant roles. Perturbation of promoters at their endogenous loci often have large phenotypic effects (Lee and Wu 2006; Yokoshi et al. 2022), whereas perturbation of redundant enhancers may only manifest their effects under challenging conditions (Frankel et al. 2010; Perry et al. 2010; Osterwalder et al. 2018). Future mutational scans of promoters and enhancers at the endogenous locus that focus on fitness effects are expected to provide insights in this direction. Secondly, different enhancers might interact with different sequence motifs in the promoter (Butler and Kadonaga 2001), which constrain promoter sequences but were not explored in the current study. Examination of a combinatorial promoter-enhancer mutation library may be required to address this possibility.

Together, ours and previous studies (Fuqua et al. 2020; Galupa et al. 2023) highlight the power of mutational scans in providing insights for developmental evolution. This approach allows us to fully explore “the possible and the actual” (Jacob 1982) of *cis*-regulatory evolution, which is currently lacking, especially in a developmental context. The differential constraints observed in different *cis*-regulatory elements can help us predict where evolutionarily relevant substitutions could occur within a locus. They also support the previous findings that the evolution of *svb* consists of multiple small-effect substitutions throughout the locus in different *Drosophila* species (Frankel et al. 2011; Preger-Ben Noon et al. 2016). In the future, mutational scans by allele replacement at the endogenous loci will provide further insights into the fitness landscape of regulatory elements in a developmental context, paralleling those in microorganisms (Metzger et al. 2015; Venkataram et al. 2016) and cell lines (Sanjana et al. 2016).

## Methods

### Promoter libraries

Random mutation libraries of *Drosophila* synthetic core promoter (DSCP), *hsp70* and *svb* promoters were synthesized at Genscript with a mutation rate of 10-20 point mutations per kb. In particular, the DSCP sequence (155 bp) was flanked by 50 bp-long sequences from *hsp70p* at each end, and the *svbp* sequence (226 bp) was flanked by 19- and 20 bp-long sequences from *hsp70p* at each end, respectively. The flanking sequences were also subjected to mutagenesis. The variants were cloned into E3N-placZattB (Fuqua et al. 2020) to replace the wild-type *hsp70* promoter, which was positioned downstream of an *E3N* enhancer and upstream of a *lacZ* reporter (Fuqua et al. 2020). The libraries were integrated into the fly genome at the attP2 site, with the injection service provided by GenetiVision. G0 transformants were crossed to w1118, and their offspring (F1) were screened for the presence of the construct by eye color. The red-eye F1 flies were individually crossed to w1118 to establish isogenic lines, which were subsequently homozygosed by sibling crosses. The mutant lines were then sequenced to identify mutations in the promoters, with primer 5’-CCAAGTTGGTGGAGTTCATAATTCC – 3’ or 5’-AGGCATTGGGTGTGAGTTCTTC - 3’. The sequences are listed in **Table S1**.

Additionally, we used two negative controls: 1) a construct with the *E3N* enhancer and *LacZ* but without any promoter; and 2) a construct with the *E3N* enhancer, a 200-bp-long “inert” spacer sequence that was computationally screened for lack of transcription factor motifs in the place of the promoter (GAAGTTTCGACTAGTCTGAAACTTCTACACAGACCGTATTAGAACTATTACTAGCTACAAGCTCCTAGTG CTTTGAAAGCTATAACCTTAAGATGCTGTTAGTATCTCAACCGACTTACTGCAGAGACTTGACGAATTCTGAAAGTTCAGAACTAGTCTCTGAGTTGCGAGGTACATTTAGCAATGTAAGAACCTCGGCT), and *LacZ*. The control constructs were integrated at attP2 sites as described above.

### Embryo collection and immunostaining

Embryos were collected from an overnight laying period at 25°C, using a standard fixation protocol (Galupa et al. 2023). During fixation and staining, a wild-type promoter control was always included in each batch, to account for batch effects.

Expression of *lacZ* was detected with a chicken anti-βGal antibody (1:500, abcam ab9361). ELAV was stained with mouse anti-ELAV supernatant (1:25, Developmental Studies Hybridoma Bank Elav-9F8A9) as a fiducial marker for the automated imaging pipeline to rotate the images, as well as for the experimenter to visually stage the embryos. For DSCP, *E3N* and *hsp70p* libraries, as well as for comparing DSCP and *svbp* activity (data in **Fig. 1, Fig. 3, Fig. S3and Fig. S5**), AlexaFluor 488 and 633 (1:500) were used as secondary antibodies for βGal and ELAV, respectively. Due to the extremely weak signal of *svbp* lines, we used extra staining steps for the *svbp* mutation library to enhance the signal (data in **Fig. 2**). After a secondary incubation of AlexaFluor 555/488 (goat anti chicken, 1:500), Biotin conjugate was used as tertiary antibody (donkey anti sheep, 1:500, 1hr incubation) and NeutrAvidin 550 was used for quaternary staining (1:500, 30min to 1hr incubation). AlexaFluor 488/647 (1:500) was used as the secondary antibody for ELAV in this case.

The stained embryos of DSCP and *svbp* libraries were mounted in ProLong Gold with DAPI. A subset of DSCP lines and all of the *hsp70p* lines were mounted in benzyl alcohol/benzyl benzoate (BABB) (Fuqua et al. 2021) and were analyzed qualitatively due to lower imaging quality. The mutation libraries were imaged on a Zeiss LSM 880 confocal microscope with an automated pipeline under a 20x objective (air, 0.8 NA) as previously described (Fuqua et al. 2021) or manually under the same setting. Embryos used for comparison between DSCP and *svb* promoter in **Fig. 1** were imaged manually under a 25x (oil, 0.8 NA) objective.

### Quantification of lacZ expression

We focused on cells in the second abdominal stripe (A2) in stage 15 embryos for analyzing the pattern and intensity of *lacZ* expression. In each embryo, the A2 region was manually selected, max-projected, and background-subtracted with a rolling ball radius of 50 pixels. To select for *LacZ*-expressing cells in the region, we first performed a Gaussian blur with a radius of 2 pixels to remove noise, and then identified regions of interest (ROIs) by automatically thresholding the image with the Otsu method in ImageJ (**Fig. S1A**). The ROIs were applied to the background-subtracted image and analyzed with the `Analyze Particles` function to extract mean intensity of each ROI. Mean intensity per embryo was calculated by 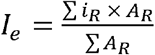, where *i* was the mean intensity and *A*_*R*_ was the area of the ROI, respectively.

The embryo mean intensity of mutant lines was compared to the wild-type in the same batch with Wilcoxon test. In the case where two biological replicates from different batches showed inconsistency in expression changes (i.e. one was different from wild-type and other was not), we took a conservative approach and removed both of them. A few *svbp* lines (57-4, 19-3, 64-14, 69-1, 89-13, 89-70, 90-7) were imaged along with two wild-type controls that were different from the control of other batches, and their intensities were scaled to the other control by linear conversion, to eliminate differences caused by background variation in the different controls. The data were normalized to the control line in each batch when combined in one plot. Data from biological replicates were merged.

To compare expression level between DSCP and *svbp*, we extracted nucleus intensity by identifying local maxima with a prominence of 2000 and selecting a circular region with a radius of 0.55 um around each local maximum. Mean intensity of the circular regions was extracted from the background-subtracted images (**Fig. S1B**).

### Allele replacement with CRISPR

We performed allele replacement following a two-step process, using a 3XP3-RFP marker as an intermediate step to easily select for integration events (Lamb et al. 2017). A 182-bp-long sequence of *svb* promoter immediately upstream of *svb* 5’UTR was targeted with two gRNAs, 5’-cgagatattcgccgttgctc-3’ and 5’-gaatacagtaagttgcgagc-3’, which were cloned into pCFD4. A repair template containing the 3XP3-RFP sequence (1.86kb) (Lamb et al. 2017) and a 983bp-long homology arm at each end was synthesized and cloned into pUC57. The gRNA (75 ng/ul) and the repair template (225 ng/ul) were mixed and injected into a fly stock expressing Cas9 in the germline (BDSC#51324: w[1118]; PBac{y[+mDint2] GFP[E.3xP3]=vas-Cas9}VK00027). Flies from the injection were crossed to an FM6 balancer and subsequently screened for RFP expression in the eyes, which indicates successful replacement of *svb* promoter by the 3XP3-RFP cassette. The RFP-positive transformants were then homozygosed for both RFP and GFP markers to establish a fly line for the second round of allele replacement.

In the second round, we replaced the 3XP3-RFP sequence with the DSCP sequence. The gRNAs were designed based on the fused sequence of *svb* locus and 3XP3-RFP cassette: 5’-GGTACCGTACGAGATCTCTC-3’ and 5’-GGCGCCTAAGGATCGATAGC-3’, cloned into pCFD4. The repair template contained a 255-bp-long DSCP sequence and the same homology arms as above. A mixture of plasmids carrying gRNAs and repair template was injected into the RFP-positive line mentioned above. Flies from the injection were crossed to an RFP/FM6 line and screened for loss of RFP. The resulting transformants were then homozygosed to establish a stable line of *svbPromoter*Δ::*DSCP* genotype. The integration was confirmed by PCR and sequencing. A list of fly strains used in this study is provided in **Table S2**.

### Fluorescent in situ hybridization

*svb* transcripts were detected with DIG-labeled probes of *svb* as per (Tsai et al. 2019). Fixed *Drosophila* embryos were mounted in ProLong Gold + DAPI mounting media (Molecular Probes, Eugene, OR) and imaged on a Zeiss LSM 880 confocal microscope with FastAiryscan under a 63x objective (Carl Zeiss Microscopy, Jena, Germany). Inside nuclei with *svb* transcription sites, the center of the transcription site was identified using the find maximum function of Fiji/ImageJ. A circle with a diameter of 12 pixels [0.85 µm, region of interest (ROI)] centered on the transcription site was then created. The integrated fluorescent intensity inside the ROI was then reported. The intensity presented in the figures is the per-pixel average intensity with the maximum readout of the sensor normalized to 255.

### Cuticle preparation

Embryos from an overnight laying period were dechorionated with bleach and left in distilled water at room temperature for 24h. After 24h, the hatched larvae were transferred onto a glass slide and mounted in Hoyer’s medium mixed with lactic acid (1:1). The slide was baked at 55°C for 2 days before being imaged with dark field microscopy.

## Supplementary Material

**Figure S1**. Quantification of *LacZ* expression in A2 cells.

**Figure S2**. Expression of *LacZ* in constructs with different promoters and without promoters.

**Figure S3**. Expression pattern of *E3N* variants.

**Figure S4**. Sequence conservation of *svb* and *eve* promoters.

**Figure S5**. Additional data for DSCP and *hsp70p* variants.

**Table S1**. Variant sequences used in this study.

**Table S2**. Fly strains used in this study.

## Acknowledgement

We thank Alessandra Reversi and Matthew Benton for providing the injection service at EMBL. We thank Noa Ottilie Borst for her help with in situ stainings, Natalia Misunou for reading the manuscript, Rafael Galupa for sharing reagents and other members of the Crocker group for their input on the project. We also thank members of Ella Preger-Ben Noon group, Alexander Stark group and Christa Buecker group for feedback on the preprint. X.C.L. is supported by a fellowship from the European Molecular Biology Laboratory Interdisciplinary Postdoc Programme (EIPOD) under Marie Skłodowska-Curie Actions COFUND (664726). Research in the Crocker lab is supported by the European Molecular Biology Laboratory (EMBL).

## Author contributions

Conceptualization: X.C.L., T.F., M.E.B., J.C. Investigation: X.C.L., T.F., M.E.B., J.C. Methodology: X.C.L., T.F., J.C. Formal analysis: X.C.L., J.C. Data curation: X.C.L. Visualization: X.C.L., J.C. Software: X.C.L., T.F., J.C. Supervision: J.C. Project administration: X.C.L., J.C. Funding acquisition: J.C. Writing, original draft: X.C.L., J.C. Writing, review & editing: X.C.L., T.F., M.E.B., J.C.

