## Supplemental figures for "Mutational scans reveal differential evolvability of *Drosophila* promoters and enhancers"

Fig. S1

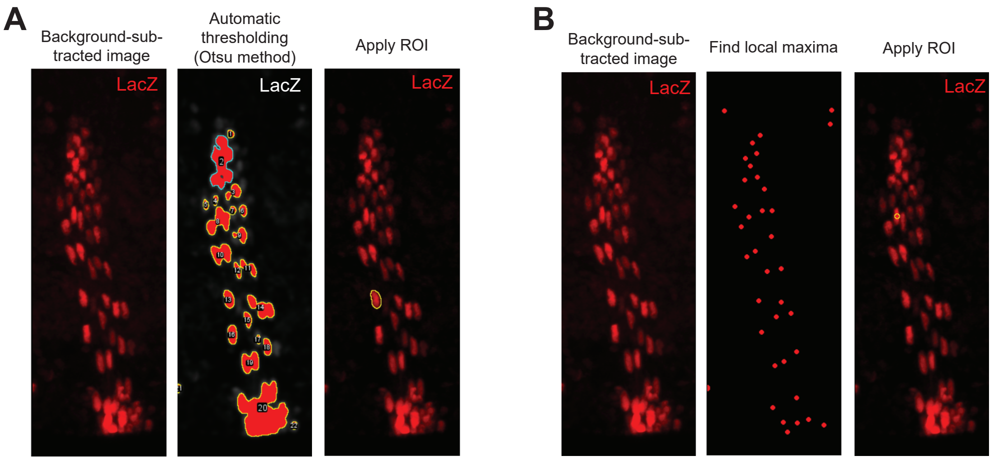


**Supplementary Figure 1. Quantification of *LacZ* expression in A2 cells.** (**A**) The background-subtracted image (left) was thresholded with the Otsu method to select regions of interest (ROIs, middle). The ROIs were applied to the background-subtracted image to extract mean intensity of the ROI (right, the yellow circle shows one ROI as an example). (**B**) Local maxima with a radius of 0.55 um were identified from the background-subtracted image as ROIs. The yellow circle in the right panel shows one representative ROI.

Fig. S2

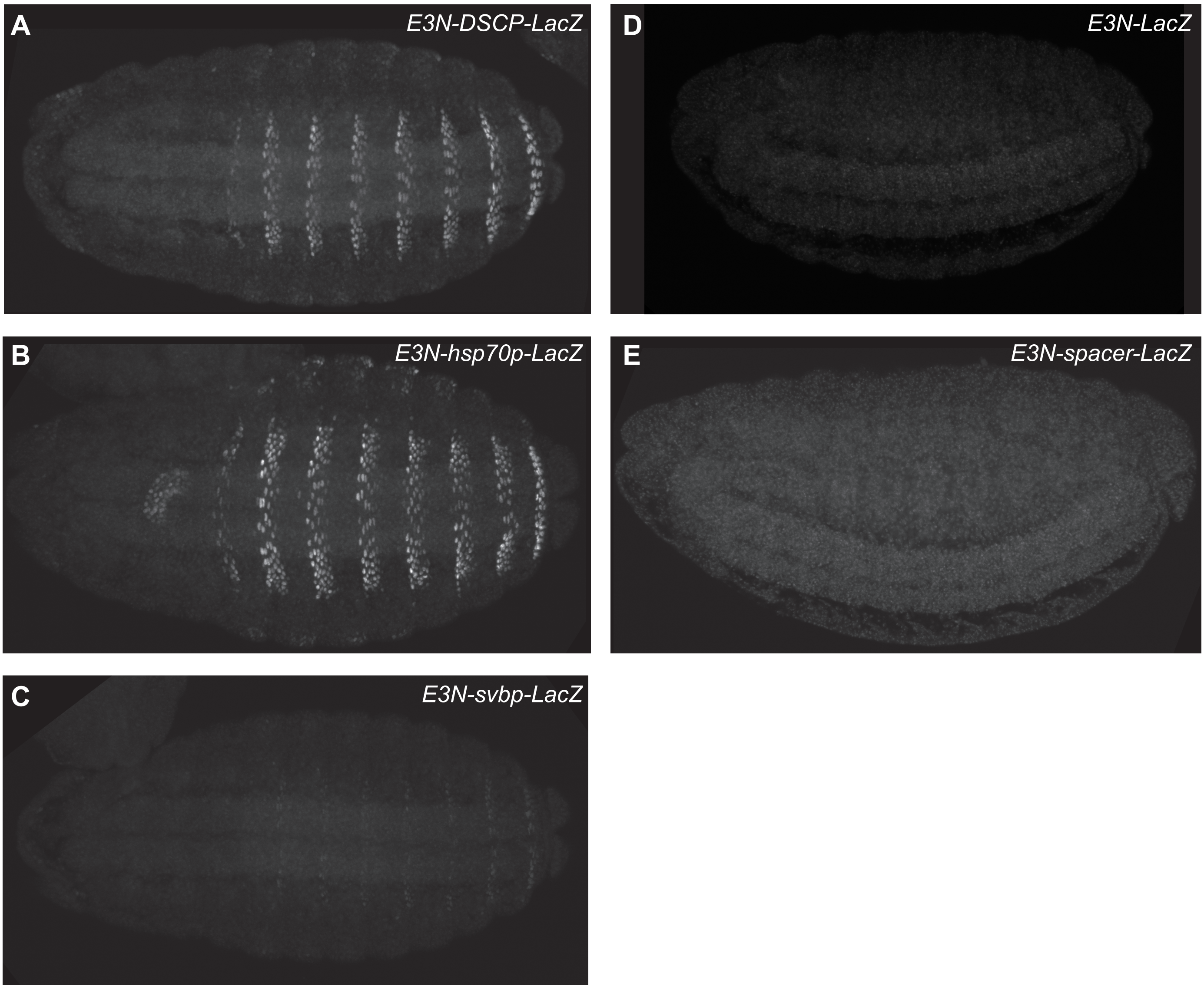


**Supplementary Figure 2. Expression of reporter constructs with different promoters and without promoters.** *LacZ* expression was detected by anti-beta-Gal immunostaining, shown in grey scale. The expression can be detected in eight abdominal stripes in stage 15 embryos for DSCP **(A)**, *hsp70p* **(B)** and *svbp* **(C)** constructs, but not in constructs without a promoter **(D)** or with a 200bp-long spacer in the place of promoter **(E)**.

Fig. S3

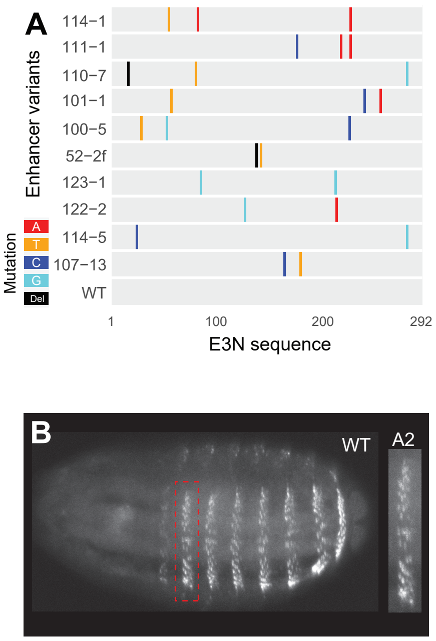


C

| Phenotype | Mutant lines |
|---|---|
| Loss of expression in most cells | 114-5 |
| Reduced to 1 column of cells | 123-1,107-13, 111-1, 110-7, 101-1 |
| Reduced to 2 columns of cells | 122-2, 114-1 |
| Sparse expression | 52-2f, 100-5 |

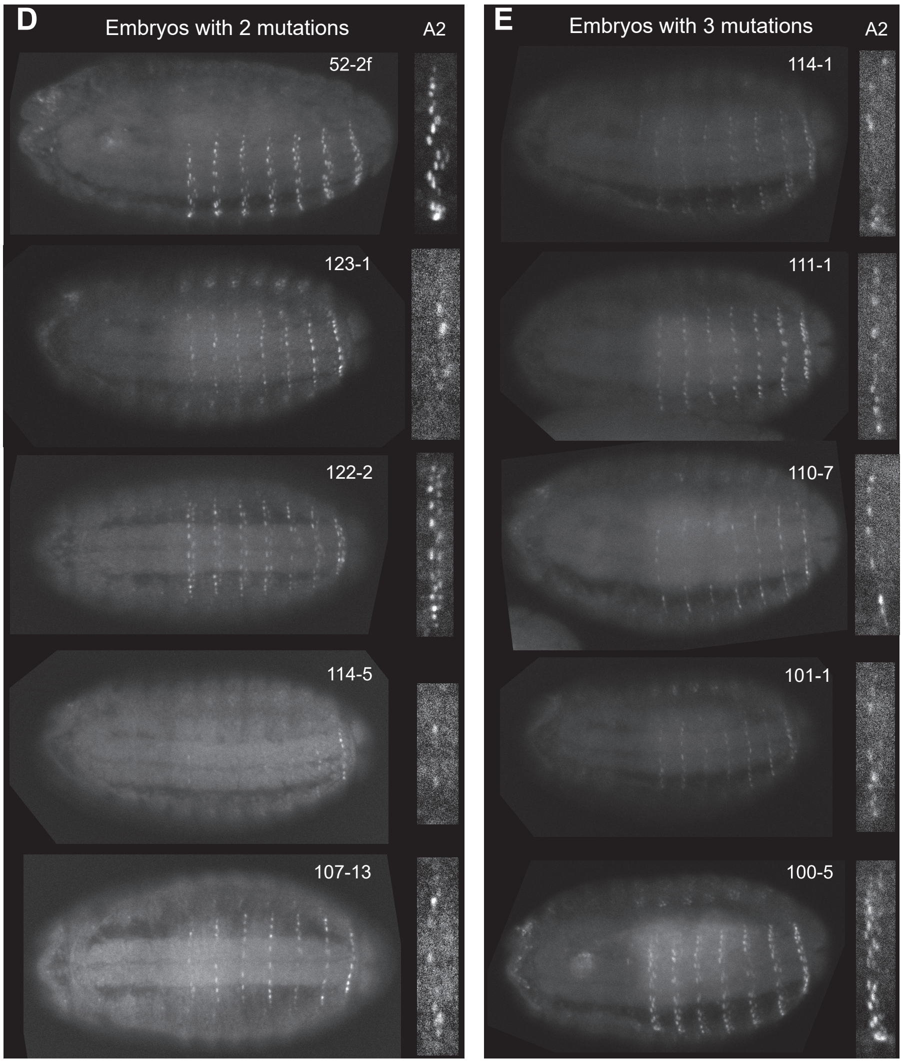


**Supplementary Figure 3. Expression pattern of *E3N* variants.** (**A**) Sequence of ten *E3N* variants sampled in this study. The mutation library was constructed by Fuqua et al. (2020), with *E3N* variants driving a *LacZ* reporter in combination with *hsp70* promoter. Del, deletion. (**B**) Wild-type pattern of *E3N* expression, showing eight stripes in the ventral abdominal region. The second abdominal stripe (A2) was selected (red rectangle, enlarged on the right) for quantification. The wild-type A2 stripe consists of 4-5 columns of cells. (**C**) Changes in the pattern of expression in *E3N* variants, classified into four catagories. Images of representative embryos are shown in (**D**) and (**E**), with lines carrying two or three mutations in *E3N*, respectively. The expression was detected by anti-beta-Gal immunostaining, shown in grey scale.

Fig. S4

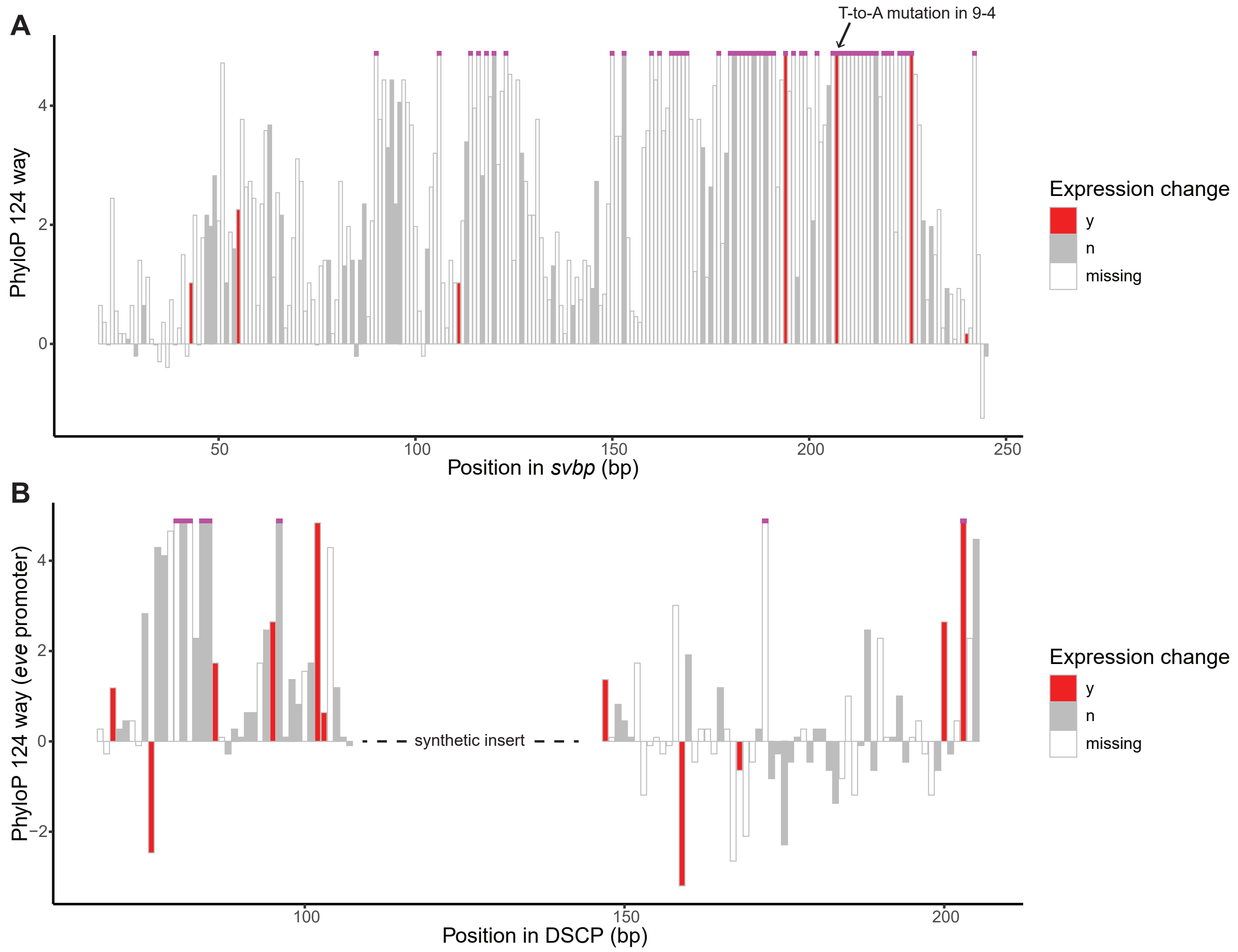


**Supplementary Figure 4. Sequence conservation of *svb* (A) and *eve* (B) promoter.** PhyloP score for 124 insects' basewise conservation was plotted for each nucleotide position in the promoters, except for the region with a synthetic insert (dash line in **B**). Scores higher than 4.88 were plotted as capped (magenta), in a similar fashion as in the UCSC genome browser. Color of the bars indicates if a mutation at a given position was only found in lines with an expression change (red) or not (grey). In the case where a position was mutated in both lines with and without an expression change, the nucleotide position was plotted as grey (i.e. the mutation was unlikely to cause an expression change when assuming additive effects). There was not a significant correlation between the presence of effects on gene expression and sequence conservation for either *svb* or *eve* promoter (Wilcoxon test, $p > 0.05$).

Fig. S5

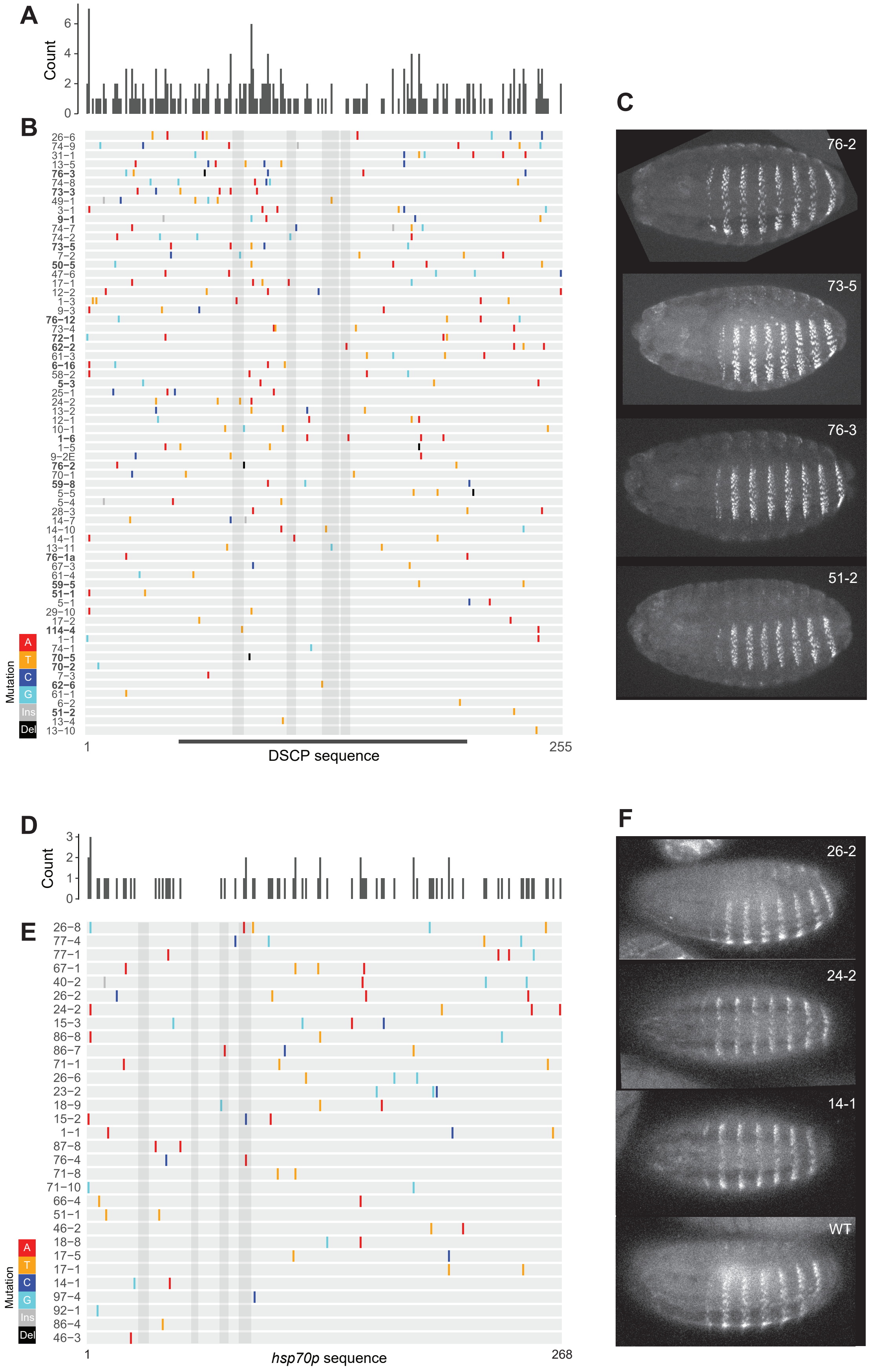


**Supplementary Figure 5. Additional data for DSCP and *hsp70p* variants.** (**A**) Distribution of mutations across 66 variants in the DSCP library, together covering 145 out of 255bp. (**B**) Sequences of DSCP variants ordered by the number of mutations, from low (bottom) to high (top). In addition to the 45 quantified variants in **Fig. 3**, this plot includes another 21 variants examined qualitatively (labels in bold). Grey shades show regions of TATA, Inr, MTE and DPE motif, respectively. (**C**) Representative images for lines examined qualitatively. (**D**) Distribution of mutations across 31 variants in the *hsp70p* library. (**E**) Sequences of *hsp70p* variants ordered by the number of mutations, from low (bottom) to high (top). Grey shades show regions of TATA, Inr, a TFIID motif and TFIID motif-DPE motif, respectively. Colored lines show the position and identity of mutations. Ins, insertion. Del, deletion. (**F**) Representative images of *hsp70p* lines. The expression was detected by anti-beta-Gal immunostaining, shown in grey scale.
